# Rnd3 regulates cell morphodynamics by spatial restriction of cell contraction signaling

**DOI:** 10.1101/2025.09.23.677979

**Authors:** Arya Sachan, Carolin Gierse, Suchet Nanda, Xiaoyi Xin, Yao-Wen Wu, Leif Dehmelt

**Affiliations:** Faculty of Chemistry and Chemical Biology, TU Dortmund, 44227 Dortmund; IMPRS for Living Matter, 44227 Dortmund; Dortmund Life Science Center (DOLCE), TU Dortmund, 44227 Dortmund; SciLifeLab and Department of Chemistry, Umeå Centre for Microbial Research, Umeå University, 90187 Umeå, Sweden

**Keywords:** Rnd3, RhoA, Rac1, PAK, Protrusion, Retraction, Contraction

## Abstract

Cell migration is enabled by dynamic changes in cell shape, which are controlled by spatio-temporal activity patterns of the Rho GTPases Rac1 and RhoA. Classical models proposed that these activity patterns are generated by mutual inhibition between the front signal Rac1 and the back signal RhoA, leading to opposing gradients that define the direction of cell migration. However, direct measurements of signal crosstalk showed that Rac1 can activate RhoA, which is incompatible with mutual inhibition. Furthermore, opposing Rac1 and RhoA gradients generated by mutual inhibition would need to overlap at least partially in the cell center. In contrast, both Rac and Rho activities were largely absent in the cell center, and both found to be highly localized near the cell edge in the cell periphery. Here, we hypothesized that these spatio-temporal Rho GTPase activity patterns are generated by the mutual inhibition between RhoA and the unconventional Rho family member Rnd3. Using rapid, optogenetic and chemical perturbations, we confirmed this mutual inhibitory crosstalk. However, we found that this crosstalk does not lead to the expected mutually exclusive spatio-temporal patterns of Rnd3 activity and cell retraction in spontaneously migrating, unperturbed cells. Instead, Rnd3 activity was even slightly elevated during cell retraction and was surprisingly depleted during cell protrusion. We discovered that this depletion is caused by the inhibition of Rnd3 by Rac1 activity, and that this newly identified inhibition is much more pronounced compared to the inhibition of Rnd3 by Rho. By combining rapid optogenetic perturbations with pharmacological manipulations, we found that Rac1 inhibition by Rnd3 is mediated by p21-activated kinases (PAKs). Investigations into the function of Rnd3 showed that it is required for the tight spatio-temporal regulation of the cell contraction/retraction signal Rho, and that it prevents ectopic, highly dynamic, spontaneous Rho activity pulses within the whole cell attachment area. Interestingly, protrusion-retraction dynamics were also severely inhibited in the absence of Rnd3, and its overexpression stimulated this process. Taken together, we show that Rac and not Rho acts as the major Rnd3 inhibitor in cells. Furthermore, our findings support a mechanism, in which Rnd3 acts as a global inhibitor of Rho that spatially restricts Rho activity to regions near the edge of migrating cells.

## Introduction

Dynamic cell shape changes play a central role in many physiological processes, for example in directional migration of immune cells or during cancer metastasis. These cell shape changes result from a combination of local cell protrusion and cell retraction, which are controlled by signal networks that operate locally in subcellular regions near the plasma membrane. The family of Rho-type small GTPases plays a central role in these signal networks (Ridley, 2015). In particular, the GTPase Rac1 stimulates cell protrusion, while the GTPase RhoA stimulates cell contraction. Classical models for the role of these central regulators in cell migration proposed that mutually inhibitory crosstalk between Rac1 and RhoA might stabilize protrusion and retraction in distinct areas of individual cells, leading to cell polarization and directional cell migration (Guilluy et al., 2011). However, recent studies from our lab could not confirm mutual inhibition between Rac1 and RhoA, but instead even showed that Rac1 can activate Rho (Nanda et al., 2023). Furthermore, these studies suggested that this crosstalk might play a role in coupling Rac and Rho dynamics to drive dynamic cell protrusion/retraction cycles at the leading edge of migrating cells. Additionally, these studies also showed that the Rac and Rho activities do not segregate into mutually exclusive front and back regions. Instead, both activities were predominantly restricted within a very thin region near the cell edge of migrating cells. Therefore, these observations suggest that Rac and Rho do not directly inhibit each other. However, inhibitory crosstalk with a third signal molecule, which is localized to more central cell attachment areas, could plausibly explain how the Rac and Rho activities are restricted near the cell edge.

Interestingly, previous studies have shown that Rho activity is mutually inhibitory with the unconventional Rho-type GTPase Rnd3 (Aoki et al., 2016). Rnd3 is a constitutively active (CA) GTPase that was shown to inhibit Rho activity by recruiting the Rho GAP p190RhoGAP (Riento & Ridley, 2006) (Wennerberg et al., 2003). Rho in turn inhibits Rnd3 via ROCK1-mediated phosphorylation at multiple sites resulting in 14-3-3 binding and translocation from the plasma membrane to the cytosol (Komander et al., 2008; Riou et al., 2013). Thus, the mutual inhibition between Rho and Rnd3 could act as a toggle switch and result in mutually exclusive activity patterns in cells. In particular, as Rnd3 is considered to be a Rho inhibitor that acts more globally on the level of the entire cell, this crosstalk might inhibit Rho activity in central cell areas and thereby play a role in restricting Rho activity near the cell edge.

Indeed, we confirmed mutual inhibition between Rho and Rnd3, however, inhibition of Rnd3 by RhoA was relatively weak. Instead, we found that active Rac1 strongly inhibits Rnd3 via a mechanism that is dependent on group I p21-activated kinases (PAKs). Investigations into the function of Rnd3 revealed that this molecule is essential to prevent ectopic Rho activity pulses throughout the whole cell attachment. Taken together, our findings show that Rnd3 acts as a global inhibitor of cell contraction dynamics, which is selectively released either by active Rac or by active Rho, thereby restricting cell contraction dynamics selectively to the dynamic edge of migrating cells.

## Results

### Mutual inhibition between Rho and Rnd3 in living cells

To directly investigate the crosstalk between Rnd3 and Rho, we combined rapid, light-based signal network perturbation methods with monitoring of the signal network response. First, to measure the effect of Rnd3 on Rho, we used the previously developed LOVTRAP system (Wang et al., 2016). This allows the manipulation of the cytosolic concentration of a protein of interest by sequestering it to the surface of mitochondria via a light-sensitive interaction. Here, we fused Rnd3 with the Zdark1 (Zdk1) domain, which interacts in the dark with a light-oxygen voltage 2 (LOV2) domain, which was targeted to the outer mitochondrial membrane. Illumination with blue light (445nm) leads to a conformational change in the LOV2 domain, which releases Zdk1-Rnd3 into the cytosol (Figure 1a), where it then can freely interact with other signal proteins of the cell.

**Figure 1.**
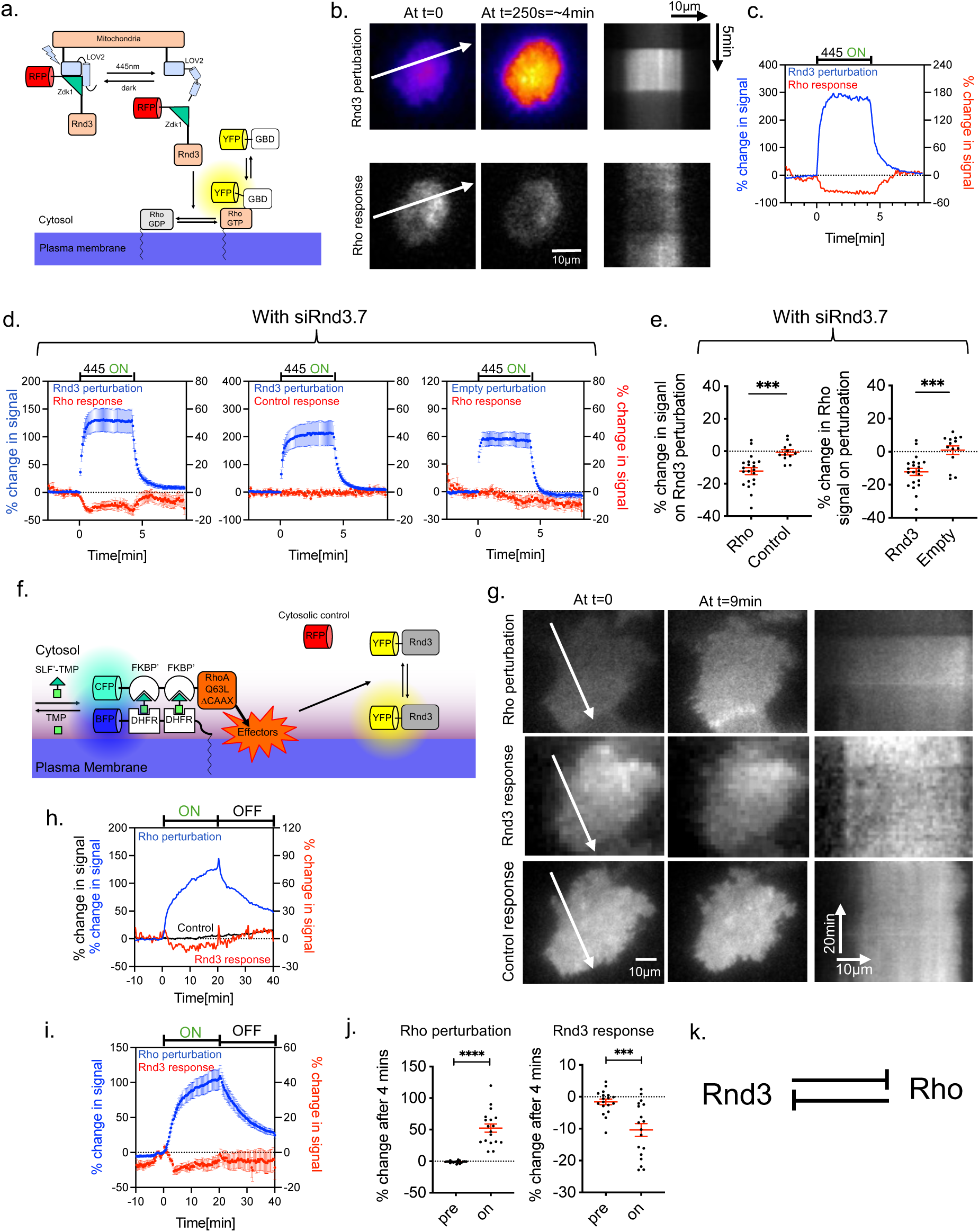
Direct investigation of mutually inhibitory crosstalk between Rho and Rnd3 activity in living cells. **a-e.** Investigation of the Rnd3/Rho activity crosstalk using the LOVTRAP system in A431 cells after knockdown of endogenous Rnd3. See Figure S1 a-b for corresponding expeirments without Rnd3 knockdown. **a.** Schematic for rapid, reversible Rnd3 perturbation via the LOVTRAP system and readout of the Rho activity sensor response. **b.** Representative TIRF images and kymographs corresponding to the white arrow in cells expressing the Rnd3 perturbation and the Rho activity sensor constructs (see also Supplementary Movie 1). **c.** Quantification of Rnd3 perturbation and parallel measurement of the Rho activity sensor recruitment dynamics in the entire cell attachment area of the representative cell shown in b. **d.** Corresponding average measurements from multiple cells along wih measurements using control sensors or an empty control perturbation. **e.** Quantification of the sensor response 1 min after Rnd3 perturbation. *n* = 3 independent experiments with >11 cells per condition. **f-j.** Investigation of the Rho/Rnd3 activity crosstalk in A431 cells. **f.** Schematic of rapid, reversible Rho GTPase activity perturbation via chemically induced dimerization and readout of the Rnd3 activity response. **g.** Representative TIRF images and kymographs corresponding to the white arrow in cells expressing the Rho perturbation, the Rnd3 and control sensor constructs. To increase signal to noise for the images and kymograph of Rnd3 measurements, the raw pixel values were downscaled by averaging 8×8 or 4×4 pixel regions, respectively All subsequent quantifications were based on the original raw pixels values without downscaling (see also Supplementary Movie 2). **h.** Quantification of Rho perturbation and parallel measurement of the uncorrected, raw Rnd3 sensor and raw control sensor response in the entire cell attachment area of the representative cell shown in g. **i.** Corresponding average measurements of the Rho perturbation and normalized Rnd3 sensor response from multiple cells. **j.** Quantification of Rho perturbation and normalized Rnd3 response before (pre) and 4 mins post-dimerizer addition (on). *n* = 3 independent experiments with 18 cells. **k.** Proposed crosstalk between Rnd3 and Rho activity in A431 cells.\*\*\*\**P* < 0.0001; \*\*\**P* < 0.001; ns: not significant; two-sided Student’s t-test. Error bars represent standard error of the mean.

We combined this rapid perturbation method with measurements of the Rho activity state at the plasma membrane via a translocation sensor based on the Rhotekin GTPase binding domain (GBD) (Benink & Bement, 2005). The Rhotekin GBD selectively interacts with the active, GTP-bound form of Rho, which is typically localized at the plasma membrane (Symons & Settleman, 2000), and not with the inactive, GDP-bound form that is sequestered in the cytosol via RhoGDIs (Be & Mccormick, 1993; Garcia-Mata et al., 2011). Specifically, we used a previously published approach (Nanda et al., 2023), in which we used a variant of this sensor that contains two tandem GBDs to increase its affinity to active Rho and that is expressed at very low levels via the delCMV promoter (Watanabe & Mitchison, 2002). To measure the local amount of active, endogenous Rho, we then used TIRF microscopy to sensitively detect the translocation of this sensor from the cytosol to the plasma membrane. Using this method, we initially didn’t observe any change in the Rho sensor signal even after the rapid release of a substantial amount of Rnd3 from mitochondria into the cytosol (Supplementary figure S1a,b). We hypothesized that endogenous Rnd3 might already be present in saturating amounts, such that this further increase of Rnd3 levels does not lead to a measurable effect. Therefore, we combined this procedure with a knockdown of endogenous Rnd3 via RNA interference. Based on western blot analysis, we were able to efficiently knockdown Rnd3 by up to 92.5 ± 3% with the most efficient siRNA (siRnd3.7, Figure S1c-d). In this condition, a rapid increase of Rnd3 lead to a significant inhibition of Rho activity (Figure 1b-e and Supplementary Movie 1) confirming the reported role of Rnd3 as a negative regulator of Rho.

Next, to measure the effect of Rho on Rnd3, we combined a chemical dimerizer-based approach to recruit active Rho to the plasma membrane with a readout of Rnd3 activity (Figure 1f). Briefly, the perturbation was based on two fusion proteins: a constitutively active mutant of RhoA (Q63L) linked to two tandem FKBP domains, and a plasma membrane anchor attached to two tandem eDHFR domains. We then induced the heterodimerization of these fusion proteins by the addition of the small molecule dimerizer SLF’-TMP, which leads to the rapid plasma membrane recruitment of active RhoA. We already established this perturbation method using these constructs in a previous study and showed that plasma membrane targeting of active RhoA efficiently induced the expected cell contraction phenotype (Nanda et al., 2023). Furthermore, plasma membrane targeting can be reversed by addition of the dimerization competitor TMP, and our previous study showed that this treatment also reversed the Rho induced cell contraction phenotype.

As an unconventional Rho-type GTPase, Rnd3 is constitutively active and thus always exists in the GTP bound state (Foster et al., 1996). Instead, the translocation of Rnd3 itself to the plasma membrane or to the cytosol is considered to be the main mechanism for switching the activity of Rnd3 on or off, respectively. We therefore measured the Rnd3 activity state simply by following the plasma membrane association of a low expressing fluorescent fusion protein (delCMV-mCitrine-Rnd3) via TIRF microscopy. With this strategy, we were able to confirm the previously published inhibitory action of Rho on Rnd3 (Figure 1g-j and Supplementary Movie 2) (Riou et al., 2013). Interestingly, while we were able to detect a rapid drop of Rnd3 quickly after Rho activation, the inhibition of Rnd3 subsequently weakened despite the continued presence of active RhoA, suggesting the presence of additional, adaptive mechanisms (Figure 1h-i).

Together, these results confirm the mutual inhibition between Rnd3 and Rho (Figure 1k), however, the inhibition of Rnd3 by Rho was relatively weak and only occurred transiently.

### Rnd3 activity dynamics in migrating cells

The previous experiments confirmed mutual inhibition between Rho and Rnd3, making them promising candidates to generate mutually exclusive activity patterns that might restrict Rho activity near the cell edge. To investigate this idea, we measured spatio-temporal Rnd3 activity patterns during spontaneously migrating A431 cells. As Rho activity is strongly enriched during cell retraction in these cells (Nanda et al., 2023), we expected that the mutual Rho/Rnd3 inhibition would lead to the local depletion of Rnd3 from such retractions. We were very surprised to observe that Rnd3 was depleted during cell protrusion rather than retraction (Figure 2a and Supplementary Movie 3). To quantitatively evaluate Rnd3 dynamics in relation to cell protrusion and retraction, we used a previously established analysis approach (Nanda et al., 2023), that is based on a modification of the ADAPT ImageJ plugin (Barry et al., 2015).

**Figure 2:**
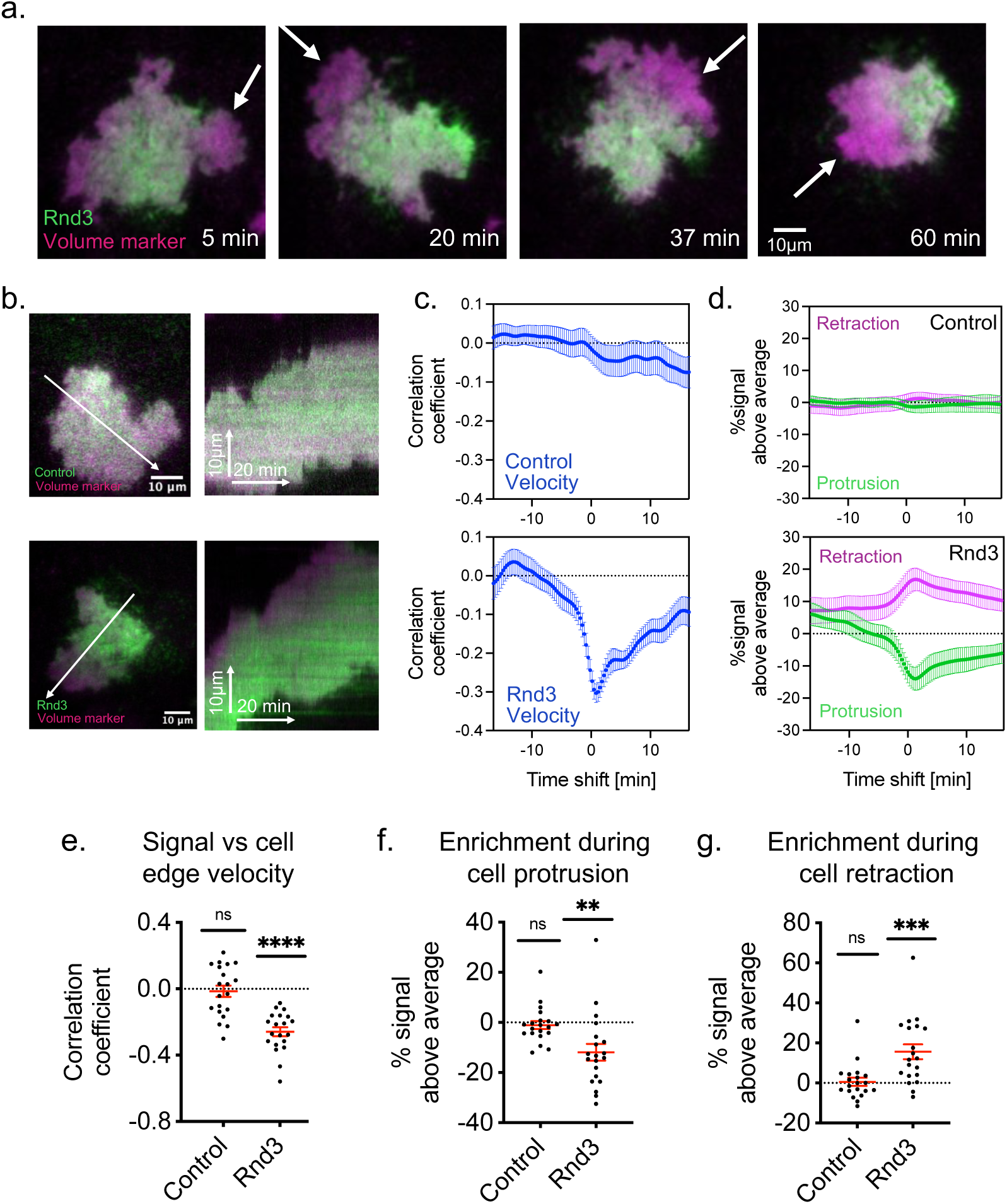
Rnd3 dynamics in migrating cells. **a.** Sequential TIRF images of a single migrating A431 cell co-expressing Rnd3 and a volume marker. Newly formed protrusions are highlighted with white arrows (see also Supplementary Movie 3). **b.** Representative TIRF images of mCitrine-Rnd3 or mCitrine as control, co-expressed with mCherry as a volume marker (left), and kymographs (right) corresponding to the white arrows in left panels (see also Supplementary Movie 4 and 5). **c.** Crosscorrelation between Rnd3 or control signals and cell edge velocity, plotted against the time shift between these measurements. **d.** Enrichment of mCitrine-Rnd3 signals in protrusions (>0.075 μm/min) and retractions (<−0.075 μm/min). Cell edge velocity was measured using the cell volume marker and all enrichment values were normalized to the average cell volume marker signal enrichment signals. **e-g.** Measurements of the signal vs cell edge velocity correlation coefficient (**e**), of the signal enrichment during cell protrusion (**f**) and of the signal enrichment during cell retraction (**g**) at a time shift of 0 min. *n* = 3 independent experiments with 20 cells per condition. \*\*\*\**P* < 0.0001; \*\*\**P* < 0.001; \*\**P* < 0.01; ns: not significant, one sample two-sided t-test. Error bars represent standard error of the mean.

These measurements clearly confirmed our qualitative observations: First, using cross-correlation analysis, we found that Rnd3 activity signals correlated negatively with cell edge velocity, with an average delay of ∼ 50 secs (Figure 2c). It should be noted that cross-correlation analysis cannot distinguish between a signal decrease during protrusion and a signal increase during retraction, since both events would result in a negative cross-correlation value. We therefore also measured the Rnd3 signal in protrusions and retractions relative to the whole cell attachement area (Nanda et al., 2023). This analysis revealed that Rnd3 is strongly depleted during cell protrusion by ∼ 20%, with a delay of about ∼ 80s after maximal cell protrusion (Figure 2d, bottom). Furthermore, Rnd3 was even slightly enriched during retraction by ∼ 10%, again with a delay of ∼ 80s (Figure 2d, bottom).

This observation suggests that Rnd3 is not only linked to cell retraction and contractile signals as suggested by several previous studies, but also might be linked to cell protrusion signals. To test, if this link between Rnd3 and cell protrusion signals is direct or indirect, we investigated the crosstalk between Rac1 and Rnd3.

### Analysis of crosstalk between Rac and Rnd3

To directly investigate the crosstalk between Rac1 and Rnd3, we again combined our rapid perturbation methods with readouts of the Rnd3 activity response. First, we used a chemical dimerizer-based approach, analogous to Figure 1f, to target a constitutively active Rac1 mutant (Q61L) to the plasma membrane by addition of SLF’-TMP and subsequently added the small molecule competitor TMP to reverse this perturbation. As shown in Figure 3a-d and Supplementary Movie 6, this dimerizer-mediated Rac1 activity perturbation leads to a substantial inhibition of Rnd3, with a rapid onset within seconds, reaching a maximum after 5-10 minutes.

**Figure 3:**
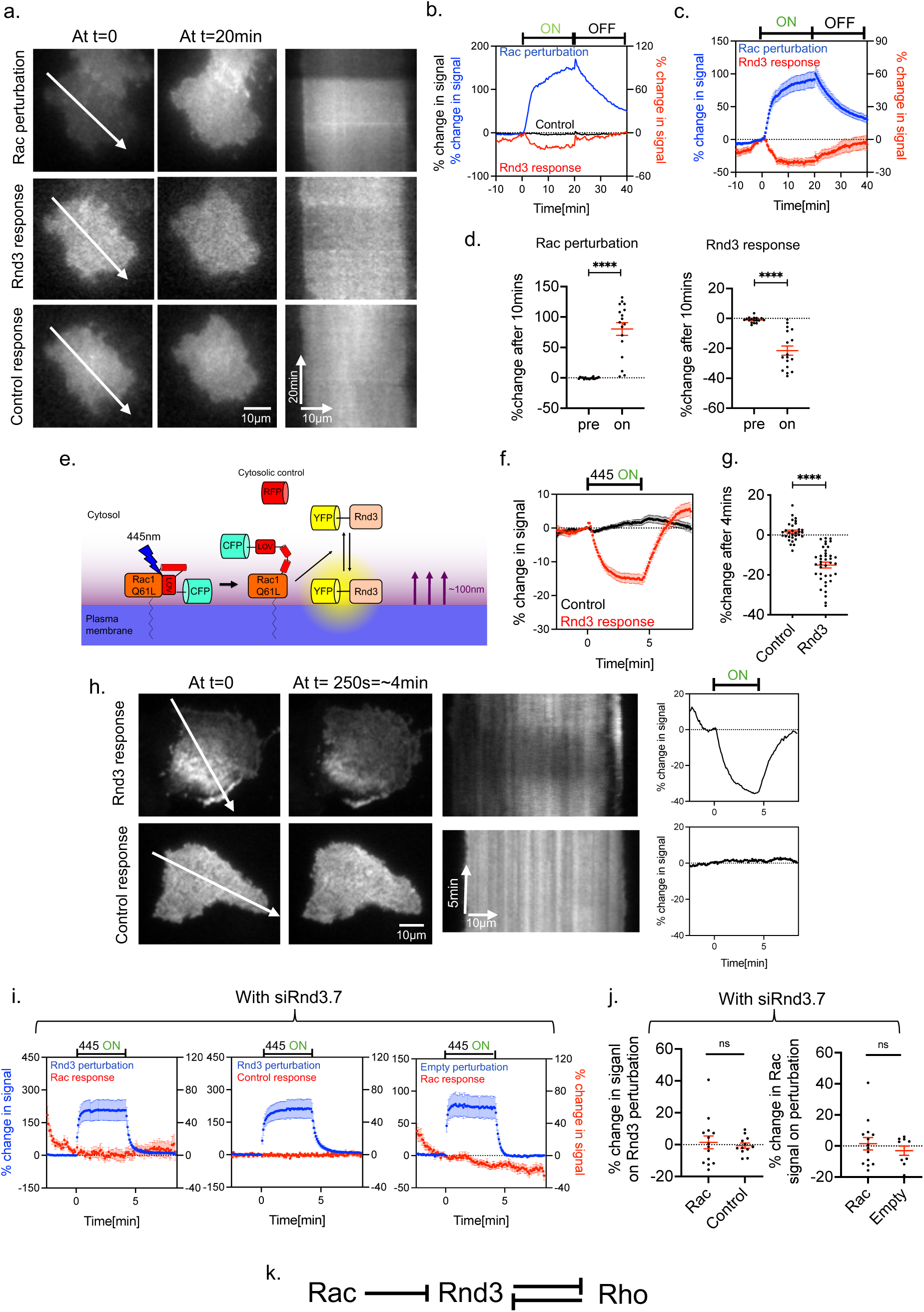
Direct investigation of activity crosstalk reveals unidirectional inhibition of Rnd3 by Rac1. **a-d.** Investigation of the Rac1/Rnd3 activity crosstalk in A431 cells via chemically-induced dimerization. **a.** Representative TIRF images and kymographs corresponding to the white arrows in cells expressing the Rho perturbation, the Rnd3 and control sensor constructs (see also Supplementary Movie 6). **b.** Quantification of Rac1 perturbation and parallel measurement of the uncorrected, raw Rnd3 sensor and raw control sensor response in the entire cell attachment area of the representative cell shown in a. **c.** Corresponding average measurements of the Rac1 perturbation and normalized Rnd3 response from multiple cells. **d.** Quantification of Rac1 perturbation and normalized Rnd3 response before (pre) and 10 min post-dimerizer addition (on). *n* = 3 independent experiments with 17 cells. **e-h.** Investigation of the Rac1/Rnd3 activity crosstalk in A431 cells via optogenetic control of photoactivatable Rac (PA-Rac1). **e.** Schematic representation of the optogenetic strategy to investigate Rnd3 recruitment by active Rac1. **f**. Quantification of the Rnd3 and control sensor response during PA-Rac1 activation in the entire cell attachment area of multiple cells. **g.** Quantification of the sensor response 4 min after photoactivation (on). *n* = 3 independent experiments with >31 cells per condition. **h.** Representative TIRF images (right panels) and kymographs (middle panels) corresponding to the white arrow in cells expressing PA-Rac1 and the Rnd3 or control sensor constructs and quantification of the Rnd3 or control sensor response in the entire cell attachment area of the corresponding representative cells shown (right panels) (see also Supplementary Movie 7). **i-j.** Investigation of the Rnd3/Rac activity crosstalk using the LOVTRAP system in A431 cells after knockdown of endogenous Rnd3. **i.** Quantification of Rnd3 perturbation and parallel measurement of the Rac activity sensor recruitment dynamics in the entire cell attachment area, including corresponding measurements using a control sensor or an empty control perturbation. **j.** Quantification of the sensor response 1 min after perturbation. *n* = 3 independent experiments with ≥9 cells per condition. **k.** Proposed model of the cellular signaling network that links Rac, Rnd3 and Rho activity in cells. \*\*\*\**P* < 0.0001; ns: not significant two-sided Student’s t-test. Error bars represent standard error of the mean.

As the observation of Rnd3 depletion during cell protrusion and after Rac1 activation was unexpected, we repeated these experiments using an alternative Rac1 perturbation method. In this second method, we used a photoactivable Rac1 construct (Wu et al., 2009), which is based on a reversible conformational change in a fusion protein of constitutively active Rac1 with the LOV2 domain. Illumination with 445 nm light is known to induce partial unfolding of the LOV2 domain. This results in the loss of an intramolecular inhibitory interaction and thereby releases the activity of the fused, constitutively active Rac1 mutant (Figure 3e). We combined this perturbation approach with Rnd3 activity measurements and confirmed that Rnd3 is rapidly and substantially inhibited by active Rac1 (Figure 3f-h and Supplementary Movie 7).

These results therefore clearly establish a direct and rapid activity crosstalk that inhibits Rnd3 downstream of Rac1 activity. The findings are consistent with our analysis of migrating cells, which indicated that Rnd3 is strongly depleted during protrusion phases in migrating cells (Figure 2d), which are typically mediated by increased Rac1 activity (Nanda et al., 2023).

Interestingly, using analogous perturbation and activity measurement methods, we observed a much stronger Rnd3 inhibition by active Rac compared to active Rho. Moreover, unlike the Rho activity induced Rnd3 inhibition (Figure 1f-j), which was only transient in the continued presence of active Rho and appeared to involve adaptive negative feedback regulation, the Rac activity induced suppression of Rnd3 was much more robust and fully reversible, suggesting that it is not actively limited by cellular adaptation mechanisms and thus might have a more prominent and robust effect in cells.

Furthermore, we also investigated the crosstalk between Rnd3 and Rac using an analgous approach as described above, using the LOVTRAP system in Rnd3 knock-down cells (Figure 1a). However, we didn’t observed any change in Rac signal on Rnd3 perturbation (Figure 3i,j). These results together show that there is a unidirectional crosstalk between Rac and Rnd3.

### Group I p21-activated kinases mediate inhibitory Rac1/Rnd3 crosstalk

We then investigated the underlying mechanism of Rac1-activity induced Rnd3 inhibition. First, since the best-characterized function of Rac is to promote actin polymerization to induce cell protrusions (Nobes & Hall, 1995), we investigated whether Rac1/Rnd3 crosstalk depends on filamentous actin. To test this, we measured the crosstalk between Rac1 and Rnd3 before and after adding 10µM latrunculin A, which is well known to inhibit actin polymerization (Coué et al., 1987). As shown in Figure 4b, latrunculin A had no effect on the inhibition of Rnd3 by Rac1, clearly showing that this crosstalk is not actin-dependent. Previous studies from our lab showed that Rac can activate Rho and Rho is well-known to inhibit Rnd3 via ROCK (Nanda et al., 2023; Riou et al., 2013). Thus, the inhibition of Rnd3 might be mediated via these potential links in a ROCK-dependent manner. However, addition of the ROCK inhibitor Y27632 (50µM) also did not have an effect (Figure 4c).

**Figure 4:**
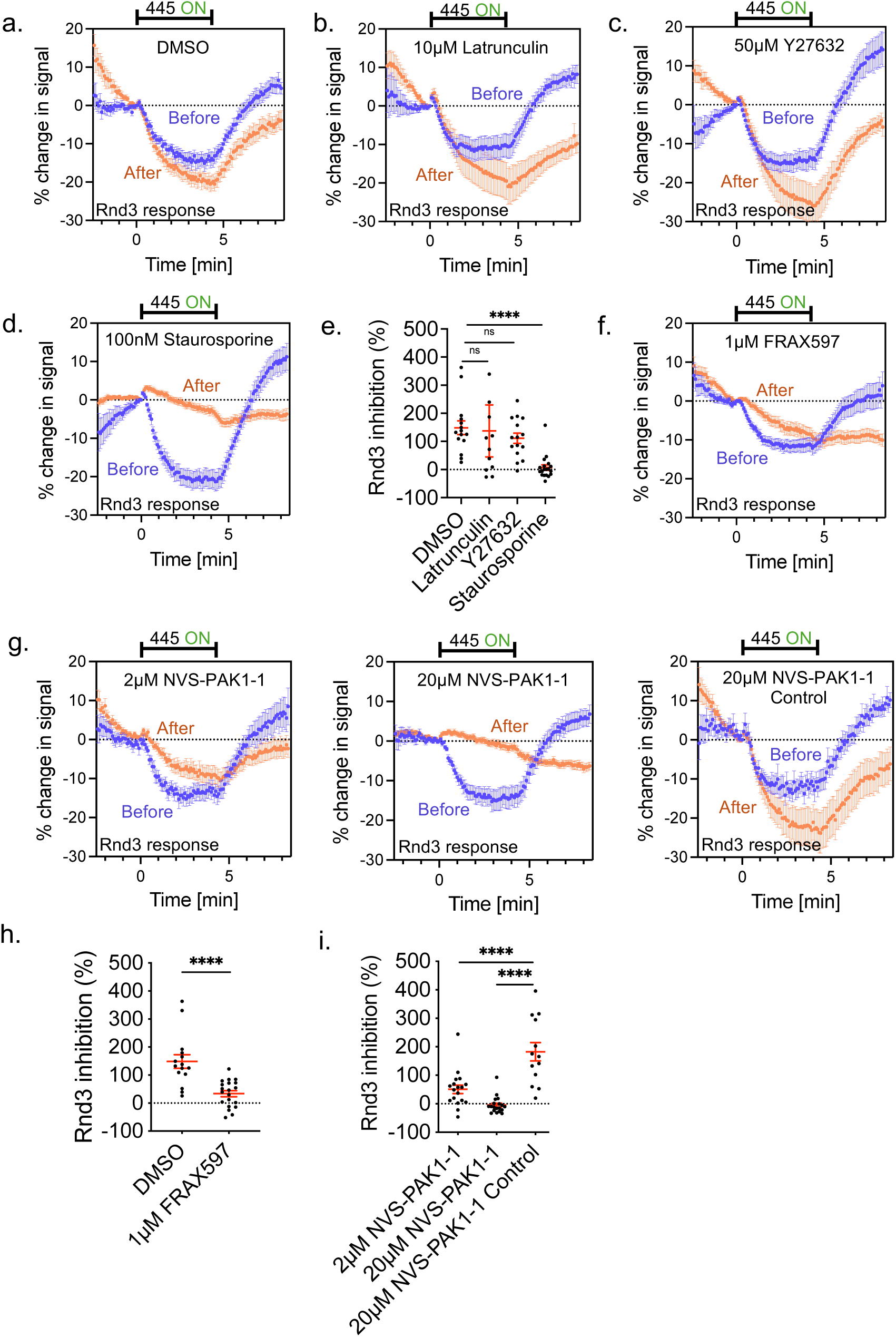
Rac1 inhibits Rnd3 via its effector kinase PAK. Measurement of Rnd3 plasma membrane recruitment during Rac1 activation in A431 cells that co-express PA-Rac1 before and 10min after adding the following small molecules inhibitors: **a.** DMSO **b.** 10µM Latrunculin**. c.** 50µM Y27632 (ROCK inhibitor) **d.** 100nM Staurosporine (pan kinase inhibitor). **f-g.** Group I PAK inhibitors: 1µM FRAX597 (**f**), or 2µM NVS-PAK1-1, 20µM NVS-PAK1-1 and 20µM NVS-PAK1-1 Control (chemically similar, inactive negative control compound) (**g**). Experiments were always performed in a paired fashion, starting with measurements before compound addition, followed by 10 minutes of compound incubation, and subsequent measurements of the same cells. **e,h,i.** Quantification of the effect of the small molecule inhibitors on Rnd3 inhibition. To quantify the effect of the inhibitors, the Rnd3 recruitment was always measured 90s after onset of Rac1 photoactivation, and presented as the percentage of the recruitment after inhibitor addition compared to the corresponding recruitment before addition. *n* = 3 independent experiments with >13 cells per condition. \*\*\*\**P* < 0.0001; ns: not significant; two-sided Student’s t-test (h), One-way ANOVA with Dunnett’s post-test using DMSO or the inacrtive NVS-PAK1-1 Control compound as negative control conditions. (e,i.). Error bars represent standard error of the mean.

We next extended our search for mediators of the Rac1/Rnd3 crosstalk and used the broad-spectrum serine/threonine/tyrosine kinase inhibitor staurosporine. Indeed, addition of 100nM staurosporine abolished the inhibition of Rnd3 by Rac1 almost to baseline levels (Figure 4d-e), clearly indicating that this crosstalk in mediated by a kinase-dependent mechanism. To further investigate which kinase is mediating this crosstalk between Rac1 and Rnd3, we investigated the role of p21-activated kinases (PAK), which are well-documented downstream effectors of Rac (Cotteret & Chernoff, 2002; Knaus & Bokoch, 1998). For this, we used two distinct group I PAK inhibitors, NVS-PAK1-1 and FRAX597, and observed that both inhibitors significantly reduced the Rnd3 inhibition by Rac1 (Figure 4f-i). At higher concentrations (20µM), NVS-PAK1-1 nearly completely abolished the Rnd3 inhibition, while a chemically similar control compound (NVS-PAK1-1 control) did not have any effect at the same concentration (Figure 4g). These results show that the crosstalk between Rac1 and Rnd3 can be disrupted by pharmacological inhibition of group I PAK kinase activity, and thereby demonstrate that this particular kinase subgroup mediates the Rac1/Rnd3 activity crosstalk.

### Rnd3 regulates cell morphology and cell morphodynamics

Previous studies only linked Rnd3 to retraction signals and thereby simply suggested that it is a Rho activity antagonist. Our new results now show that Rnd3 is also linked to protrusion signals. This unexpected finding suggests that Rnd3 might not just simply be a Rho activity inhibitor, but that it might play a more fundamental role in the spatio-temporal coordination between protrusion and retraction signals and thereby might function as a critical regulator of cell morphodynamics. We therefore reduced the expression of endogenous Rnd3 using RNA interference and investigated how this affects cell morphology, morphodynamics and the spatio-temporal organization of the underlying signal network.

We were able to efficiently knockdown Rnd3 via two independent siRNAs, which resulted in the reduction of protein levels by 92.5 ± 3% (siRnd3.7) and 72.05 ± 1.5% (siRnd3.6; see Supplementary figure S1c-d). First, we investigated the effect of Rnd3 knockdown on cell morphology and F-actin distribution. We expected that loss of Rnd3 would lead to higher Rho activity and thereby increased contractile activity, resulting in smaller and rounder cells that contain locally enriched F-actin structures due to the activation of F-actin-rich contractile structures. Using microscopy-based morphometric analyses, we indeed found, that the loss of Rnd3 leads to a dramatic reduction in cell adhesion area compared to control (Figure 5a,b), as well as to significantly higher maximal F-actin intensity signals in cells (Figure 5c). These observations thereby further support the already well-established inhibitory action of Rnd3 on Rho and cell contraction signaling in our A431 cell system.

**Figure 5:**
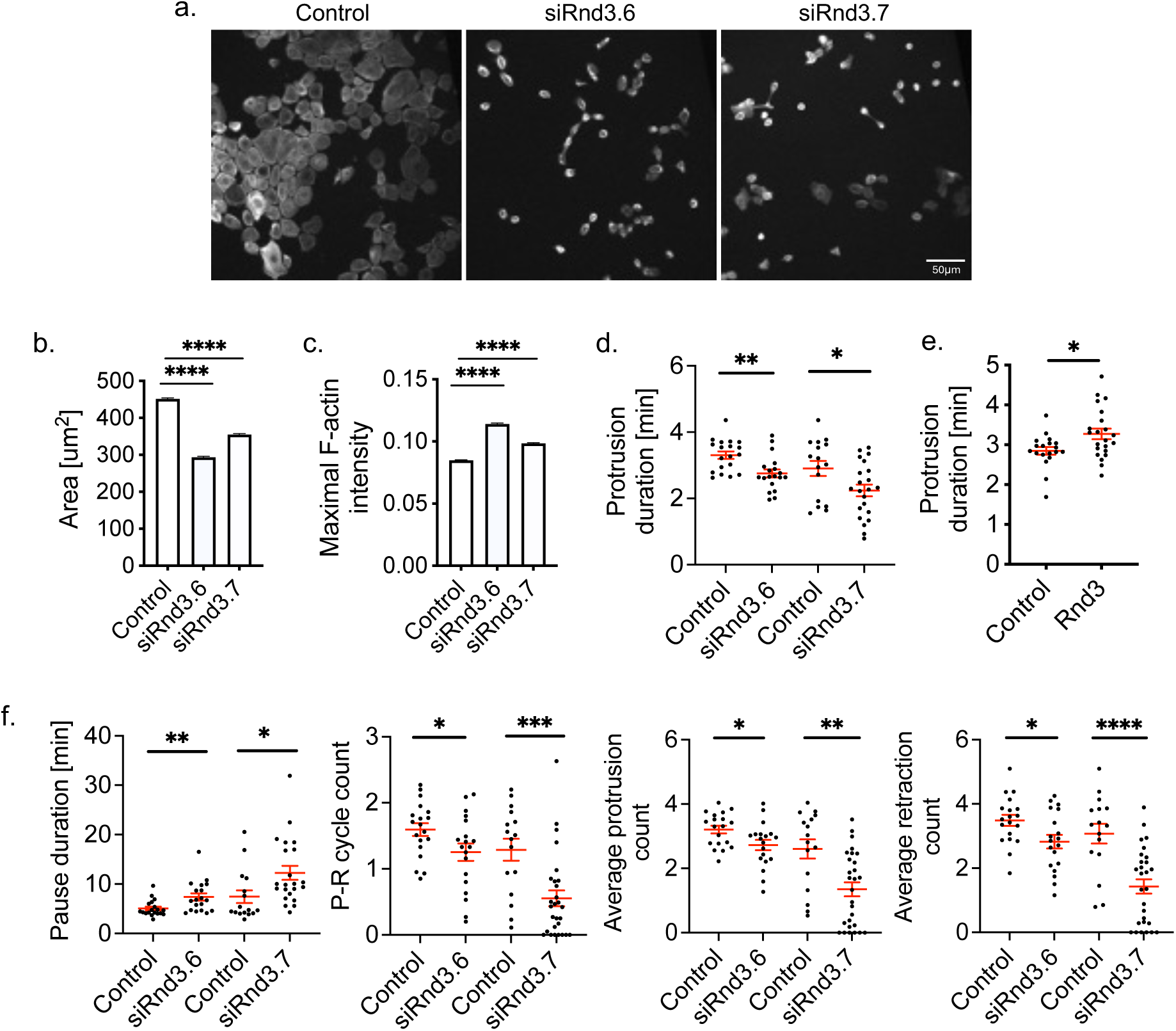
Rnd3 is a critical regulator of cell morphodynamics. **a-c.** Effect of Rnd3 knockdown on cell attachment and actin organization. *n* = 3 independent experiments with >11,420 cells per condition. **a**. Representative images of F-actin, stained via rhodamin-phalloidin, in control and Rnd3 knockdown cells. **b.-c.** Quantification of cell attachment area (b) and maximal F-actin intensity (c). Area and maximal F-actin intensity were quantified using Cellprolifer. **d-f.** Quantification of cell morphodynamics based on cell edge velocity measurements, after Rnd3 knockdown (**d,f;** *n* = 3 independent experiments with >16 cells per condition) or Rnd3 overexpression (**e;** *n* = 3 independent experiments with >19 cells per condition).\*\*\*\**P* < 0.0001; \*\*\**P* < 0.001; \*\**P* < 0.01, \**P* < 0.05; two-sided Student’s t-test (d,e,f). One-way ANOVA using Dunnett’s post test (b,c). Error bars represent standard error of the mean. Control siRNA experiments were performed independently for siRNA #6 and #7 using the same non-targeting sequence (d,f).

We also expected that loss of Rnd3 would lead to increased, global cell contraction/retraction, and allow only very short protrusions that are immediately countered by retraction. Conversely, overexpression of Rnd3 is expected to lead to decreased retraction, enabling the prolonged formation of protrusions. Indeed, measurements of cell morphodynamics showed that knockdown of Rnd3 reduced the protrusion duration in cells (Figure 5d), and overexpression of Rnd3 lead to a significant increase in the duration of individual protrusion events (Figure 5e).

As our investigations uncovered a link between Rnd3 and the cell protrusion signal Rac1, we expected that Rnd3 might also play a role in the interplay between cell protrusion and retraction. Indeed, knockdown of Rnd3 lead to a reduction in the number of protrusion-retraction cycles (Figure 5f), and changes in several additional morphodynamic parameters indicating that cells are generally less active in the absence of Rnd3 compared to control cells (Figure 5f).

### Rnd3 constrains cell retraction signals in space and time

The A431 cells that we used in our study are highly motile, and they usually do not generate a stable, polarized shape with a constant protrusive front and contractile back to migrate in a directional manner. Instead, they generate highly dynamic local cycles of protrusion and retraction to migrate in a more exploratory, stochastic fashion (Nanda et al., 2023). In these cycles, local Rac1 activity was tightly associated with local cell protrusion, which quickly switched to local Rho activity associated with cell retraction (Nanda et al., 2023). Interestingly, these activities all occur sequentially in the same local area of the leading cell edge (Figure 6a, yellow arrows, and Supplementary Movie 8). While our previous work has identified a direct crosstalk mechanism that can link Rac1 and Rho activities, it was unclear, why both Rac and Rho are nearly exclusively activated near the cell edge. This is particularly surprising, as Rho is known to amplify its own activity via positive feedback, which can lead to spontaneous pulsatory Rho activity dynamics in the entire cell attachment area in adherent mammalian cells (Graessl et al., 2017; Kamps et al., 2020).

**Figure 6:**
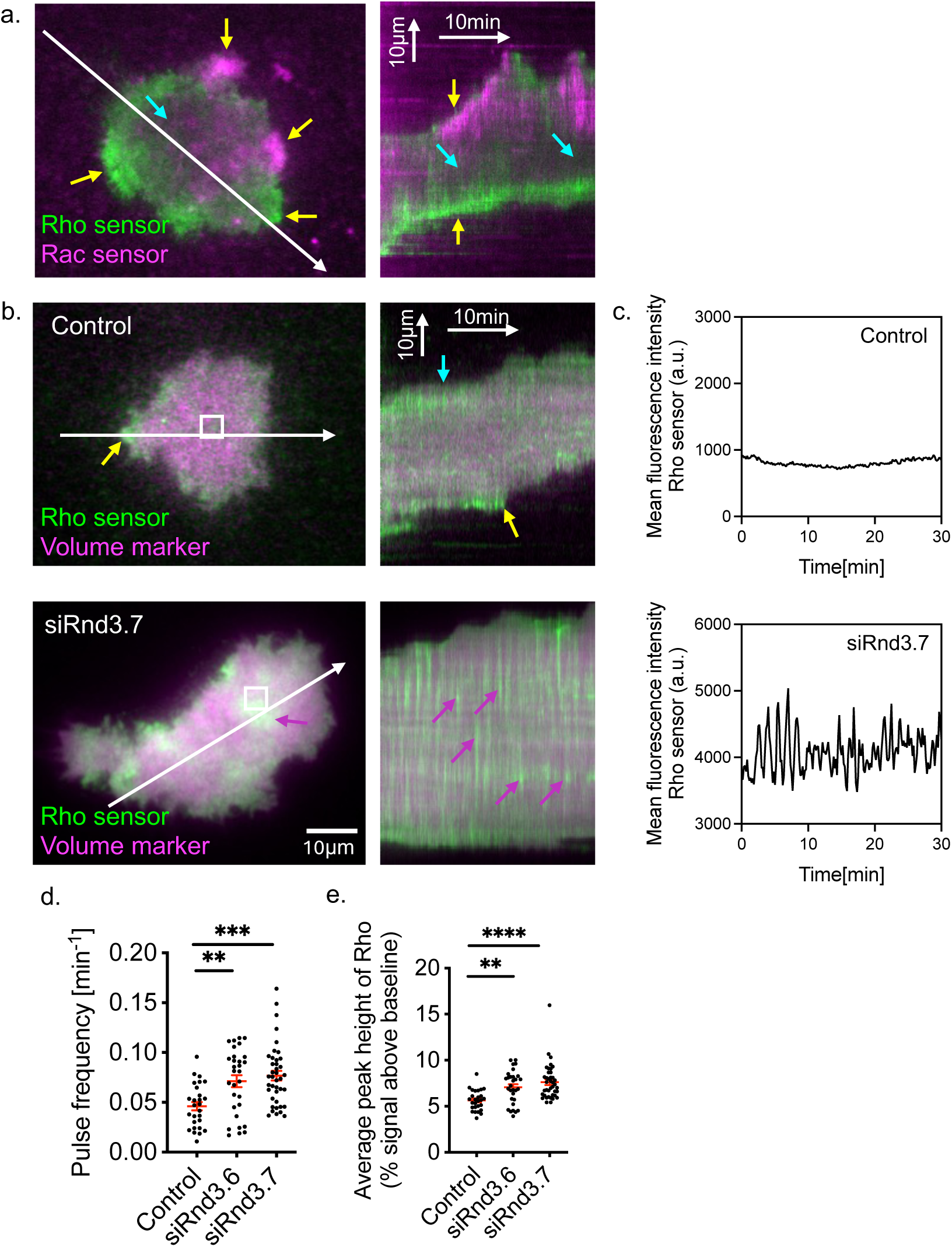
Rnd3 constrains retraction signals in cells. **a.** Representative TIRF image (left), and kymograph (right) corresponding to the white arrows of a cell co-expressing Rac and Rho activity sensors (see also Supplementary Movie 8). Yellow arrows point to increased Rac or Rho activity in peripheral cell areas, and Cyan arrows point to regions that are largely devoid of either Rac or Rho activity. **b.** Representative TIRF images (left), and kymographs (right) corresponding to the white arrows of control (top) and Rnd3 knockdown (bottom) cells, expressing Rho activity sensors and mCitrine as a volume marker (see also Supplementary Movie 9 and 10). Yellow and cyan arrows point to regions of Rho activity that are associated with cell edge retractions or pauses, respectively, and magenta arrows point to transient Rho activity pulses in central cell attachment areas. **c.** Mean fluorescence intensity of the Rho sensor corresponding to the white box in control and Rnd3 knockdown cells on the left. **d-e.** Measurement of frequency (**d**) and average peak height (**e**) of Rho activity pulses in control and Rnd3 knockdown cells. *n* = 3 independent experiment with >26 cells per condition. \*\*\*\**P* < 0.0001; \*\*\**P* < 0.001; \*\**P* < 0.01; One-way ANOVA with Dunnett’s post-test. Error bars represent standard error of the mean.

We hypothesized that Rnd3 might play a role in this process by spatially restricting contractile signals in the majority of the cell attachment area, and that its loss downstream of active Rac enables contractile signals only within regions that recently generated a cell protrusion. Therefore, to gain a better understanding of the nature of contractile Rho activity signals in the presence or absence of Rnd3, we investigated how the depletion of Rnd3 affects Rho activity dynamics in cells.

As shown in Figure 6b (top) and Supplementary Movie 9, control cells generate contractile Rho activity signals predominantly near the cell edge during retractions (yellow arrows) and during pauses (cyan arrows). In contrast, Rho activity was largely absent in central cell attachment areas. After Rnd3 knockdown, we observed a striking shift in this spatio-temporal Rho activity pattern towards central cell attachment areas, in which prominent, highly dynamic Rho activity pulses were observed (magenta arrows) (Figure 6b, bottom and Supplementary Movie 10). Quantification of these Rho activity dynamics in control and Rnd3 knockdown cells clearly showed that both, the pulse frequency and the average peak height of pulses were significantly increased after Rnd3 knockdown (Figure 6d,e). These results therefore support our hypothesis, that Rnd3 acts as a global inhibitor of Rho that spacially restricts Rho activity to regions near the edge of migrating cells.

## Discussion

Classical models of Rho GTPase signaling in cell migration proposed mutual inhibition between the cell protrusion signal Rac1 and the cell retraction signal Rho (Figure 7a, Top) (Guilluy et al., 2011). Such a crosstalk mechanism would be able to define a stable front-back polarization and the direction of cell migration by generating opposing Rac and Rho gradients that partially overlap in the cell center (Figure 7a top). However, in A431 cells, which migrate in a highly dynamic, and more exploratory fashion, we were not able to observe such stable front-back signal gradients. Instead, we observed very transient phases of Rac or Rho activity that were associated very locally with dynamic cell edge protrusion and retraction events, and both Rac and Rho activities were largely absent in central cell attachment areas (Figure 6a and Figure 7a, Bottom). We hypothesized that this absence of Rac and Rho activity might be due to another signal molecule that acts as a mutual inhibitor for Rac, Rho, or both. Rnd3 was previously proposed to be a mutual inhibitor for Rho and we confirmed this crosstalk in A431 cells (Figure 7b, Top), Furthermore, we discovered that Rnd3 depletion leads to a loss of the tight spatio-temporal control of the cell contraction/retraction signal Rho (Figure 7b, Bottom).

**Figure 7:**
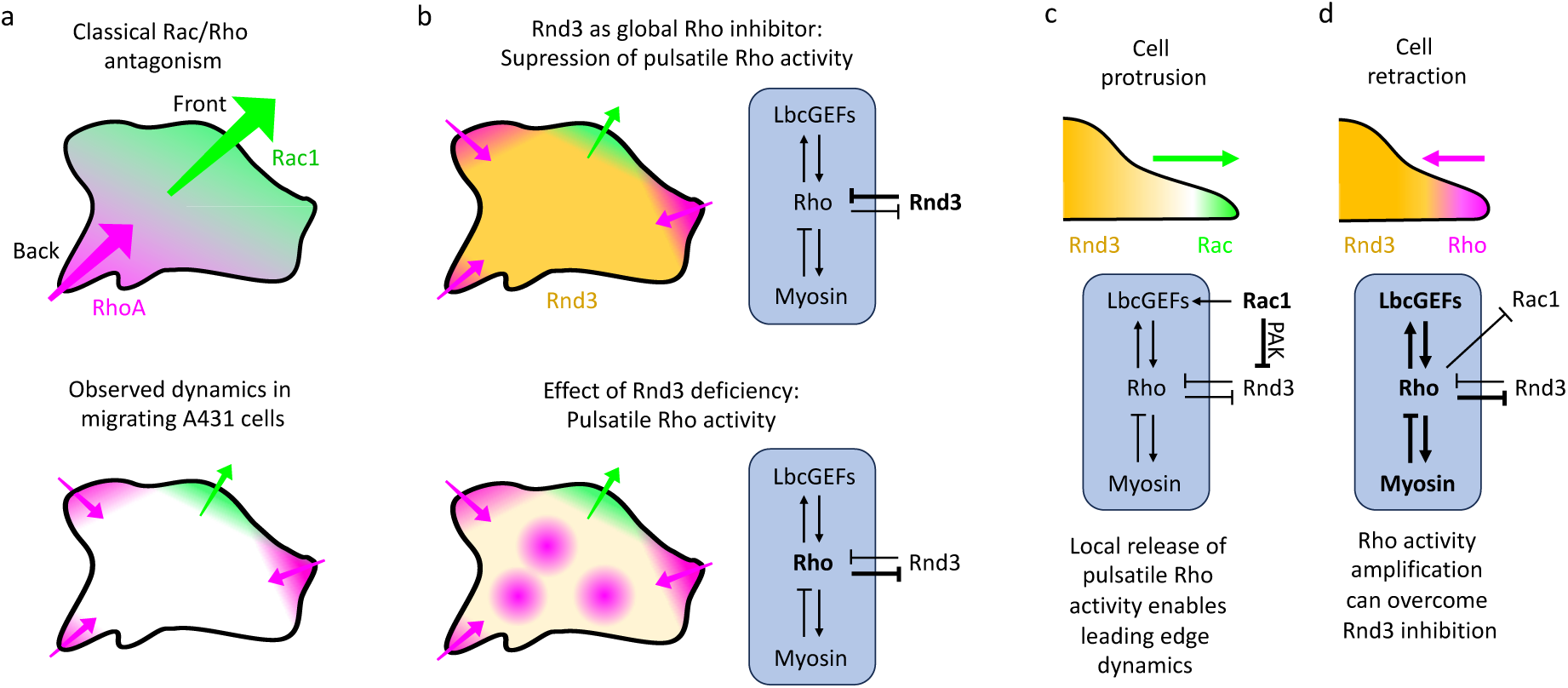
Proposed mechanism for spatial restriction of cell contraction signals by Rnd3. a-d: Schematics of spatio-temporal Rac, Rho and Rnd3 activity patterns in cells (a-b: top views; c-d: side views of the leading edge), and signal network topologies that describe causal relationships. **a.** Top: Based on the classical mechanism of mutual inhibition, cells are expected to generate mutually exclusive Rac and Rho activity gradients that partially overlap in central cell areas. Bottom: In migrating A431 cells, non-overlapping Rac and Rho activity patterns are observed in peripheral areas near the cell edge. More central cell attachment areas are devoid of either Rac or Rho signals. **b.** Top: Rnd3 pattern in A431 cells suggest a role as a global Rho inhibitor, which is only reduced in the presence of active Rac. Bottom: Rnd3 deficiency after knockdown enables pulsatile Rho activity dynamics throughout the entire cell attachment area. Pulsatile Rho activity can be explained by coupled fast positive and slow negative feedback loops (blue boxes; (Graessl et al., 2017)), and high Rnd3 levels can suppress these dynamics by inhibiting Rho. **c.** Local Rac activity in cell protrusions can release the inhibitory action of Rnd3 on Rho. This can lead to subsequent retraction, enabling leading edge protrusion/retraction dynamics. Rac can further stimulate this process by activating a subset of Lbc-type GEFs (eg. Ahrgef11/12 (Nanda et al., 2023)). **d.** Once Rho is activated, its activity can be amplified via Lbc-type GEFs (e.g. ARHGEF2 or Arhgef12 (Graessl et al., 2017)) to overcome the inhibition by Rnd3. Active Rho can inhibit Rac during retraction (Nanda et al., 2023) and delayed negative feedback inhibition of Rho via myosin can prepare cells for the next cycle (Graessl et al., 2017).

In particular, in the absence of Rnd3, the tight restriction of Rho activity near the cell edge is lost and instead, cells generate ectopic, highly dynamic Rho activity pulses within the entire cell attachment area. In our previous work, we found that such pulses can be generated by a combination of a fast positive feedback, in which Rho activators of the Lbc-GEF family and Rho amplify each other, coupled to slow negative feedback regulation, in which Lbc-GEFs are inhibited by a myosin-dependent process (Figure 7b) (Graessl et al., 2017; Kamps et al., 2020; Kowalczyk et al., 2022; Lee et al., 2010; Gierse et al., 2025;). Endogenous Rnd3 levels seem to be sufficient to effectively suppress such pulsatory dynamics in A431 cells (Figure 7b, Bottom), However, other cells, such as non-migratory, sessile U2OS osteosarcoma cells can spontaneously generate robust Rho activity pulses throughout their cell attachment area under control conditions (Graessl et al., 2017). Rac activity during cell protrusion can remove Rnd3 locally from the plasma membrane at the leading edge of the cell, thereby eliminating its inhibitory action on Rho activity specifically at these sites (Figure 7c) (Figure 2a). Conceptually, Rac1 can thus indirectly activate Rho by inhibiting its inhibitor Rnd3 (Figure 7c). We described Rac1-dependent activation of Rho already in a previous study (Nanda et al., 2023), and found that this crosstalk can also be mediated by the stimulation of the Lbc-GEF family members ARHGEF11 (PDZ-RhoGEF) and ARHGEF12 (LARG), downstream of active Rac1. Once Rho is sufficiently activated, it can amplify its activity via Lbc-GEFs, inhibit Rac (Guilluy et al., 2011; Kuo et al., 2011; Nanda et al., 2023; Ohta et al., 2006; Sanz-Moreno et al., 2008; Vicente-Manzanares et al., 2011) and stimulate myosin-based cell retraction (Figure 7d). Together, this sequence of events can generate a highly dynamic cycle of cell protrusion and retraction.

Interestingly, we do not observe a detectable reduction of Rnd3 during retraction, suggesting that Rho can somehow overcome the inhibitory action of Rnd3 during this process. We could envision two different explanations for this observation: 1) The positive feedback amplification of Rho activity during retraction is more effective than the inhibitory action of Rnd3. 2) Although Rnd3 is still present at the plasma membrane, its ability to inhibit Rho is somehow reduced, for example by a reduced ability to recruit inhibitory Rho GAPs (Wennerberg et al., 2003). Our results indeed support the idea, that Rho activity can overcome Rnd3 inhibition, which we found to be relatively weak (Figure 1f-j).

The mechanism described above puts Rac1 in a very strong position to dictate, where Rho activity dynamics are enabled, raising the question, how active Rac1 by itself is confined to the very leading edge of cells. Interestingly, we did not detect a substantial inhibition of Rac activity by Rnd3, suggesting that other mechanisms exist to prevent spurious Rac1 activity. Indeed, positive feedback between Rac1 activity and membrane curvature inducing molecules was proposed, supporting the idea that Rac1 activity might only be efficiently generated at the very leading edge of the cell (Galic et al., 2014). Thus, highly local, amplified Rac1 and cell curvature signals might be an important initial trigger that can then stimulate highly local, amplified Rho activity that is spatially constrained by Rnd3.

In conclusion, we reveal a novel inhibitory crosstalk mechanism that links the protrusion regulator Rac1 and the contraction inhibitor Rnd3, and propose a mechanism, in which Rnd3 acts as a global inhibitor that ensures that Rho stimulated cell contraction is confined to the edge of migrating cells.

## Materials and methods

### Cell culture

A431 cells (CRL-1555, ATCC) were maintained at 37°C and 5% CO2 using standard cell culture techniques in DMEM medium (10% FBS, PAN Biotech or Sigma-Aldrich Chemie GmbH/Merck and 2 mM L-Glutamine, PAN Biotech). For imaging, cells were either plated onto glass-bottom dishes (MatTek) or LabTek glass surface slides (Thermo Fischer Scientific). The glass surfaces were pre-coated with 10 μg/ml Fibronectin (45 min at RT for LabTek and overnight at 4°C for MatTek). For migration experiments, cells were plated on 35 mm culture dishes and then replated on fibronectin-coated MatTek glass-bottom dishes on the day of imaging. Lipofectamine^TM^ 3000 (Invitrogen, Thermo Fisher Scientific) was used for transfection of plasmid DNA.

### Plasmid constructs

TagBFP-2xeDHFR-CAAX for targeting proteins of interest to the plasma membrane via chemically-induced dimerization was previously described in (Liu et al., 2014). mTurquoise2-NES-2xFKBP’-RacQ61LΔCAAX, mTurquoise2-NES-2xFKBP’-RhoAQ63LΔCAAX for targeting Rac and Rho to plasma membrane via chemically induced dimerization and the control constructs delCMV-mcherry and delCMV-mcitrine were previously described in (Nanda et al., 2023). To generate CMV-mCitrine-Rnd3, the fluorophore was removed from CMV-EGFP-Rnd3 (gift from Channing Der, Addgene plasmid #23229; http://n2t.net/addgene:23229; RRID:Addgene_23229) by digestion using BspEI and AgeI-HF and combined via Gibson Assembly with an insert that was amplified using delCMV-mCitrine-RBD (Graessl et al., 2017) as the template and primers 5’-CGTCAGATCCGCTAGCGCTACCGGTCGCCACCATGGTGAGCAAG-3’ and 5’-TTCATGTCGACGGATCTGAGTCCACTTCCAGAACCGGAACCTCCGGACTTGTACAGCTCG-3’. To generate delCMV-mCitrine-Rnd3, the promoter was exchanged by ligation-based subcloning, using CMV-mcitrine-Rnd3 and delCMV-mcitrine and AseI and NheI-HF.

The PA-Rac construct (mCerulean-PA-Rac1Q61L) and pTriEx-mCherry-ZdkI were a kind gift from Klaus Hahn, University of North Carolina (Wu et al., 2009; Wang et al., 2016). pTriEx-NTOM20-CCL-moxBFP-CCL-LOV2 was previously described (Kamps et al., 2020). To generate the Rho activity sensor (delCMV-mCitrine-2xRBD), the fluorophoe was exchanged by ligation-based subcloning using delCMV-mcherry-2xRBD (Nanda et al., 2023), delCMV-mcitrine and BsrgI and AgeI. To generate the Rac activity sensor (delCMV-mCitrine-3xp67Phox), the fluorophoe was exchanged by an analogous ligation-based subcloning using delCMV-mcherry-3xp67Phox (Nanda et al., 2023), delCMV-mcitrine and BsrgI and AgeI. To generate pTriEX-mCherry-Zdk1-Rnd3 (Rnd3 perturbation construct), the small GTPase insert was removed from pTriEX-mCherry-Zdk1-RhoA Q63L (Addgene #81058) by digestion using HindIII-HF and KpnI-HF and combined via Gibson Assembly with an insert that was amplified using CMV-mcitrine-Rnd3 as the template and primers 5’-GGTTCTGGTTCTGGTGGTACCGGTTCCGGTTCTGGAAGTG −3’ and 5’-TATACAGCTGTGCGGCCGCAAGCTTTCACATCACAGTGCAGCTC −3’. The siRnd3.7 resistant Rnd3 perturbation construct (pTriEX-mCherry-Zdk1-Rnd3-siRnd3.7r) was generated by removing a part of the Rnd3 sequence in pTriEX-mCherry-Zdk1-Rnd3 by digestion using BspHI and SacI-HF and combined via Gibson Assembly with an insert that was amplified using pTriEX-mCherry-Zdk1-Rnd3 as a template and the mutagenic primers 5’-CAGATGTTAGTACATTAGTAGAACTGTCCAATCACAGGCAGACG −3’ and 5’-GTCAGATGCTCAAGGGGCTTCATGATGTCCCCATAATTTTTGGC −3’.

### siRNA mediated knockdown

Knockdown experiments were performed using ON-TARGETplus siRNAs (Dharmacon^TM^, Horizon Discovery): (siControl: #2 5′-UGGUUUACAUGUUGUGUGA-3’, Rnd3: #6 5’-CUACAGUGUUUGAGAAUUA-3’, #7 5’-UAGUAGAGCUCUCCAAUCA-3’). For knockdown, A431 cells were plated on 35mm culture dishes and transfected with 30 nM siRNA using Lipofectamine^TM^ RNAiMAX (Invitrogen). Cells were then incubated for 24h or 48h before splitting and reseeding on glass bottom dishes. For live cell experiments, the siRNA-treated cells were again transfected using Lipofectamine 3000 for expressing the plasmid constructs. Experimental analysis of knockdown cells and quantification of knockdown efficiency via Western Blots were performed 36-48h after siRNA transfection.

### Western blot analysis

Cells were washed once with ice cold PBS and subsequently lysed in ice cold 1× cell lysis buffer (Cell Signaling Technology, #9803) for 5 min on ice. Lysates were then centrifuged at 13000 × *g* for 10 min at 4 °C to remove insoluble debris. The protein concentration in the supernatant was quantified using the Bradford assay. Protein samples were prepared in 4× Laemmli buffer, boiled at 95 °C for 5 minutes, and resolved by SDS-PAGE (Bio-Rad, #4561084). Proteins were then transferred to PVDF membrane (Merck) using wet blot transfer. PVDF Membranes were briefly soaked in methanol before setting up the transfer. Transfer was performed at 70 V for 3hr at 4 °C.

Membranes were blocked using Intercept® blocking buffer (LI-COR, #927-60001) for 1 hour at room temperature. Primary antibody incubation was performed overnight at 4 °C using the following antibodies diluted 1:1000 in blocking buffer: Rnd3 (Sigma-Aldrich 05-723, Lot #3821703) and GAPDH (Cell Signaling Technology, #2118; Lot #14) as a loading control. After washing with TBS-T, membranes were incubated with fluorescently labelled secondary antibodies (IRDye 680RD anti-Mouse IgG, #926-68070, Lot #D10901-15; IRDye 800CW anti-Rabbit IgG, #926-32211, Lot #D10831-15) diluted 1:1000 in blocking buffer for 1 hour at room temperature. Following final washes, blots were visualized using the Odyssey® CLx imaging system (LI-COR).

### Microscopy

Total internal reflection fluorescence (TIRF) microscopy was carried out on an Olympus IX-81 system equipped with a motorized TIRF-MITICO illumination combiner, a high numerical aperture Apo TIRF 60×/1.45 NA oil immersion objective, and a ZDC autofocus unit. Spectral imaging for fluorophores emitting in blue (TagBFP), cyan (mTurquoise2/mCerulean), yellow (mCitrine), and red (mCherry) channels utilized a quadrupule bandpass dichroic mirror (ZT405-440/514/561) along with specific emission filters (HC 435/40, HC 472/30, HC 542/27, and HC 629/53). Excitation was achieved using either a 514 nm OBIS diode laser (150 mW, Coherent Inc., USA), or Cell R diode lasers (Olympus) at 405 nm (50 mW), 445 nm (50 mW), and 561 nm (100 mW). Additional wide-field illumination was provided by the Spectra X light engine (Lumencor). Fluorescence detection was performed using an EMCCD camera (C9100-13, Hamamatsu, Germany) operated at medium gain with no pixel binning. The imaging system was equipped with a temperature-controlled incubation chamber. Live-cell time-lapse microscopy was conducted at 37 °C at specified frame intervals in CO₂-independent, HEPES-buffered imaging medium (Pan Biotech) supplemented with 10% fetal bovine serum (FBS). For morphometric analysis, automated scanning was performed on the same microscope using wide-field illumination, a UPlanSApo 10×/0.4 NA air objective, and software-based autofocus.

### Rapid Rnd3 perturbation via the LOVTRAP system

We used the LOVTRAP system to control the cytosolic concentration of a protein-of-interest in living cells as described previously (Wang et al., 2016). Here, we used this approach to control the cytosolic concentration of Rnd3. Cells were transfected with pTriEx-NTOM20-GS-moxBFP-cc-GS-LOV2 (described previously in (Kamps et al., 2020)), pTriEX-mCherry-Zdk1-Rnd3 and delCMV-mCitrine-2xRBD (Rho activity sensor) or delCMV-mCitrine-3xp67Phox (Rac activity sensor). In experiments that were combined with Rnd3 knockdown, pTriEX-mCherry-Zdk1-Rnd3-siRnd3.7r (perturbation construct resistant to siRnd3.7) was used instead of pTriEX-mCherry-Zdk1-Rnd3. The perturbation of Rnd3 was performed using illumination with a 445nm laser in wide-field mode. Due to the high sensitivity of the LOV2 domain, we used a 100x neutral density filter (with laser intensity set to 100%) to release Rnd3 fused to Zdk1 into the cytosol. The 445 nm laser was continuously on in the perturbation time interval, except during the few milliseconds in which images were aquired. To avoid crosstalk between optogenetic control and fluorescent readouts during measurements, imaging was only conducted using the 514 nm and 561 nm laser excitation lines. BFP fluorescence was only recorded after the completion of the time series to avoid interference with optogenetic control.

The RhoGTPase activity sensor measurement *A_Rho_* was calculated by measuring the fluorescence intensity of the Rho GTPase activity sensor *I_Rho_* in the entire cell adhesion area. These raw intensity measurements were normalized by subtracting the background signal outside the cell area *I_Rho,BG_*, and dividing by the initial, background-corrected intensity value *I_Rho_*_,0_ − *I_Rho,_*_0,*BG*_ just before the perturbation.

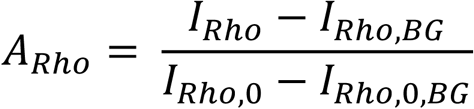

### Rapid Rac1/Rho perturbation via chemically induced dimerization

Chemically induced dimerization was performed as described previously (Liu et al., 2014). The SLF’-TMP (dimerizer) and the competitor TMP were synthesized and purified using the protocols described in (Liu et al., 2014). Cells were transfected with TagBFP-2xeDHFR-CAAX and the perturbation construct (mTurquoise2-NES-2xFKBP’-RhoAQ63LΔCAAX or mTurquoise2-NES-2xFKBP’-RacQ61LΔCAAX). To investigate Rho GTPase crosstalk, cells were co-transfected with delCMV-mCitrine-Rnd3 along with the control construct delCMV-mCherry. Dimerization was chemically induced by adding 10 μl SLF’-TMP dimerizer after 10 min and was stopped 20 min after addition of dimerizer by adding 10 μl TMP competitor.

The subsequent analysis was performed as described previously (Nanda et al., 2023). Briefly, the fluorescence intensity of Rnd3 *A_Rnd3_* was measured via TIRF microscopy in the whole cell attachment area during the entire observation period. The raw Rnd3 fluorescence intensity measurement (*I_Rnd3_*) was normalized by subtracting the background signals outside the cell area (*I_Rnd3,BG_)* and dividing by the initial, background-corrected intensity measurements before the perturbation (*I_Rnd3_*_,0_ − *I_Rnd3_*_,0,*BG*_). The corrected Rnd3 sensor measurements are then obtained by subtracting the normalized control sensor measurements from the normalized Rnd3 measurements.

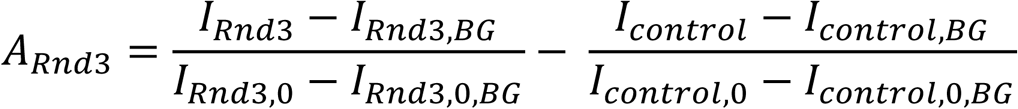

### Rapid Rac1 perturbation via PA-Rac1

The development of photo-activable Rac1 was described in (Wu et al., 2009). Here, optogenetic perturbation using this construct were performed as described previously (Nanda et al., 2023). Cells were co-transfected with mCerulean-PA-Rac1 and delCMV-mcitrine-Rnd3 and light-based activation of PA-Rac1 was performed using TIRF illumination with a 445 nm laser. To avoid oversaturation of PA-Rac1 activation, a 10000x neutral-density filter was used and the light intensity of the 445 nm laser was set to 30% to further reduce the intensity. During the photoactivation, the 445 nm laser was continuously on, except during the acquisition of images. To avoid crosstalk between optogenetic control and fluorescent readouts during measurements, imaging was only conducted using the 514 nm and 561 nm laser excitation lines. mCerulean fluorescence was only recorded after the completion of the time series to avoid interference with optogenetic control. In these experiments, delCMV-mCherry was used as a control sensor, which was either co-expressed with the Rnd3 sensor, or expressed in a separate cell population.

The subsequent analysis was performed as described previously (Nanda et al., 2023). The Rnd3 activity sensor measurement *A_Rnd3_* was calculated by measuring the fluorescence intensity of the Rnd3 activity sensor *I_Rnd3_* in the entire cell adhesion area via TIRF microscopy. These raw intensity measurements were normalized by subtracting the background signal outside the cell area *I_Rnd3,BG_*, and dividing by the initial, background-corrected intensity value *I_Rnd3_*_,0_ − *I_Rnd3_*_,0,*BG*_ just before the perturbation.

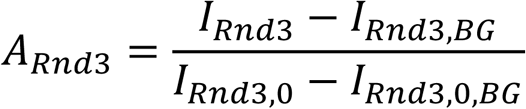

In cases where control sensor is co-expressed with Rnd3, the corrected Rnd3 activity sensor measurements were obtained by subtracting the normalized control sensor (delCMV-mcherry) intensities from the normalized Rnd3 sensor measurements.

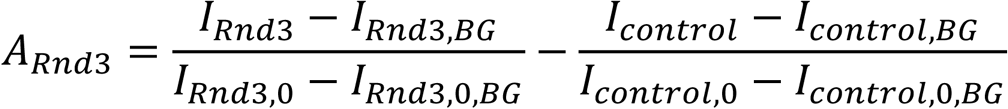

### Analysis of cell morphodynamics and local fluorescence signals at the cell edge

Analysis of cell edge velocity crosscorrelation, signal enrichment in protrusions/retractions, and quantitative analysis of cell edge movements were done as described previously (Nanda et al., 2023). Briefly, the local enrichment of Rnd3 activity was measured in A431 cells by transfecting delCMV-mCitrine-Rnd3 along the cell volume marker, delCMV-mCherry. Cell edge movements were measured based on the cell volume marker irrespective of dynamic Rnd3 activity signals using a modified version of the ADAPT plugin (Barry et al., 2015). The outputs of the ADAPT plugin were further analyzed using custom ImageJ scripts to extract velocity crosscorrelation, protrusion/retraction enrichment measurements and time intervals that correspond to protrusions, retractions and pauses. To measure the effect of overexpression of Rnd3 on cell morphodynamics, CMV-mcitrine-Rnd3 was transfected instead of delCMV-mcitrine-Rnd3. Cell edge movements > 0.075 µm/min were defined as protrusions, and < −0.075 µm/min were defined as retractions. To study the effect of siRNA mediated knockdown of Rnd3 on cell morphodynamics, delCMV-mCherry was used as cell volume marker to identify the cell borders.

### Analysis of cell morphology

Cell morphology was analysed as described previously (Lüttig et al., 2025). 24hr after siRNA transfection, cells were reseeded on fibronectin coated LabTek dishes. 8h after reseeding, the cells were washed with PBS and were fixed using prewarmed formaldehyde. The cells were then permeabilized with 0.1 % Triton X-100 and stained for F-actin and nuclei using Rhodamin Phalloidin (1:1000) and Hoechst 33342 for 30 min at RT. Imaging was performed by automated scanning of individual wells and the images were analyzed using Cell Profiler (Carpenter et al., 2006) to measure the cell attachment area and maximal F-actin intensity.

### Analysis of pulsatile Rho activity dynamics

To analyze pulsatile Rho activity dynamics, cells were initially plated on 35mm dishes, co-transfected with the Rho activity sensor (delCMV-mCherry-2xRBD) and the cell volume marker delCMV-mCitrine. On the next day, cells were reseeded on glass bottom dishes, followed by imaging. To quantify pulsatile Rho activity dynamics, a custom-made ImageJ script that was described previously (Graessl et al., 2017) was used with minor changes.

### Software for image and video analysis

All the video and Image analysis was performed using ImageJ (https://imagej.net/software/fiji/). The analysis of kymographs was performed using the built-in multi kymograph ImageJ plugin. Parameters of migrating cells were analysed using a modified version of the ADAPT plugin as described above in more detail (Barry et al., 2015; Nanda et al., 2023). Cell Profiler (Carpenter et al., 2006) was used to quantify maximal F-actin intensity and cell adhesion area, and data were further processed using MatLab. Statistical analyses and data plotting were performed using Prism 10 (GraphPad).

## Author Contributions

L.D. and A.S. designed the research. A.S. performed, analyzed, and optimized the majority of experiments. C.G. initiated the project and supported the data analysis. S.N. performed parallel Rac and Rho sensor mesurements in individual cells. Y.W.W. and X.X. synthesized the chemical dimerizer. L.D. supervised experiments. L.D. and A.S. wrote the majority of the manuscript. All authors contributed to discussions and manuscript preparation.

## Acknowledgements

We thank Sven Müller (MPI Dortmund) for expert microscopy support and the late Philippe Bastiaens (MPI Dortmund) for departmental support. We also would like to thank Ricarda Lüttig for support in quantifying cell morphology and helpful discussions. We would like to thank Dr. Sonja Sievers from COMAS – Compound Management and Screening Center, MPI Dortmund for providing the PAK inhibitor, NVS-PAK1-1 and NVS-PAK1-1_Control. This work was supported by the Deutsche Forschungsgemeinschaft DFG project grant 823/9-1 and DFG Principal Investigator grant DE 823/10-1 to L.D., a Doctoral research grant from the German Academic Exchange Service (DAAD) to A.S., the European Research Council, ERC (ChemBioAP) to Y.W.W., the Knut and Alice Wallenberg Foundation to Y.W.W., the Göran Gustafsson Foundation for Research in Natural Sciences and Medicine to Y.W.W. and Vetenskapsrådet (Nr. 2018-04585, Nr. 2022-02932) to Y.W.W.

