## Supplementary Information for "Rnd3 regulates cell morphodynamics by spatial restriction of cell contraction signaling"

### Supplementary Figure:

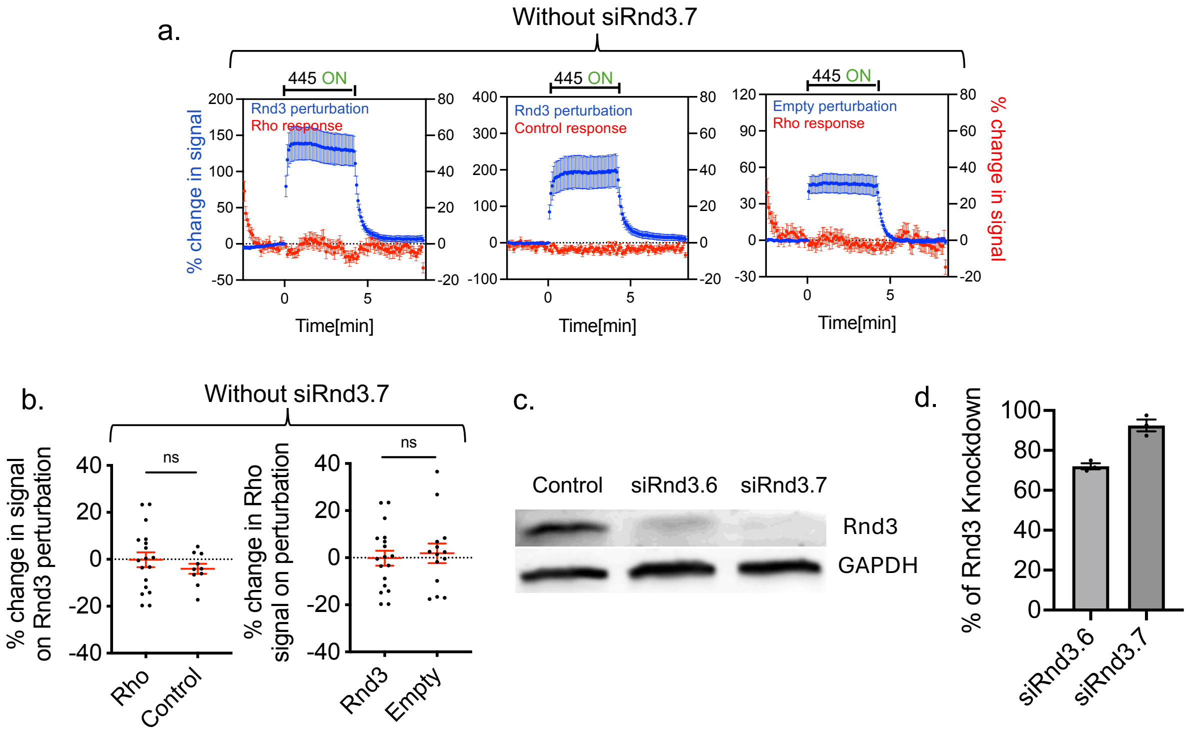


**FigureS1. a-b.** Investigation of the Rnd3/Rho activity crosstalk using the LOVTRAP system in A431 cells in the presence of endogenous Rnd3. See Figure1 a-e for corresponding expeirments in combination with Rnd3 knockdown. **a.** Quantification of Rnd3 perturbation and parallel measurement of the Rho activity sensor recruitment dynamics in the entire cell attachment area from multiple cells along wih measurements using control sensors or an empty control perturbation. **b.** Quantification of the sensor response 1 min after Rnd3 perturbation**.** $n = 3$ independent experiments with ≥10 cells per condition. **c-d** Analysis of knockdown efficiency via Western blot analysis 36h after siRNA transfection. **c.** Representative western blot image confirming the knockdown of Rnd3. **d.** Quantification of knockdwon efficiency via densitometry. ns: not significant; two-sided Student's t-test.

### Description of Supplementary Movies:

Supplementary Movie 1: Measurement of Rnd3/Rho crosstalk in A431 cells in the entire cell attachment area (related to Fig. 1b). Time-lapse TIRF videos of Rnd3 perturbation in Rnd3 knockdown A431 cells and parallel Rho activity response. Photouncaging of Rnd3 was performed by 445 nm laser illumination. Images were collected with a frame rate of 12/min.

Supplementary Movie 2: Investigation of RhoA/Rnd3 crosstalk in A431 cells in the entire cell attachment area via chemically-induced dimerization (related to Fig. 1g). Time-lapse TIRF videos of the Rho perturbation and the control and Rnd3 response. Perturbation of Rho was induced via the chemical dimerizer SLF'-TMP and reversed using the competitor TMP. Images were collected with a frame rate of 3/min. To reduce noise, 8x8 pixels of the raw data were scaled down to a single pixel using averaging.

Supplementary Movie 3 and 5: Spatio-temporal activity patterns of Rnd3 aquired together with a volume marker (related to Fig. 2a and 2b (bottom)). Time-lapse TIRF video of Rnd3 (delCMV-mCitrine-Rnd3) and the volume marker (delCMV-mCherry) in an A431 cell. Images were collected with a frame rate of 3/min.

Supplementary Movie 4: Spatio-temporal activity patterns of a corresponding control construct aquired together with a volume marker (related to Fig. 2b (top)). Time-lapse TIRF video of the control construct (delCMV-mCitrine) and the volume marker (delCMV-mCherry) in an A431 cell. Images were collected with a frame rate of 3/min.

Supplementary Movie 6: Investigation of Rac1/Rnd3 crosstalk in A431 cells in the entire cell attachment area via chemically-induced dimerization (related to Fig. 3a). Time-lapse TIRF videos of the Rac perturbation and the control and Rnd3 response. Perturbation of Rac was induced via a chemical dimerizer SLF'-TMP and reversed using the competitor TMP. Images were collected with a frame rate of 3/min.

Supplementary Movie 7: Measurement of Rac1/Rnd3 crosstalk in A431 cells in the entire cell attachment area using photoactivable Rac1 (related to Fig. 3h). Time-lapse TIRF video of the Rnd3 activity and control response in A431 cells expressing photoactivatable Rac (PA-Rac1), which was activated using 445nm light. Images were collected with a frame rate of 12/min.

Supplementary Movie 8: Spatio-temporal activity patterns of the Rac and Rho activity sensors in an A431 cell (related to Fig. 6a). Time-lapse TIRF video of the Rac (delCMV-mCherry-3xp67Phox) and Rho (delCMV-mCitrine-2xRBD) activity sensors. Images were collected with a frame rate of 6/min.

Supplementary Movie 9: Spatio-temporal activity patterns of the Rho activity sensor with a volume marker in a control A431 cell (related to Fig. 6b (top)). Time-lapse TIRF video of the Rho activity sensor (delCMV-mCherry-2xRBD) and volume marker (delCMV-mcitrine). Images were collected with a frame rate of 6/min.

Supplementary Movie 10: Spatio-temporal activity patterns of the Rho activity sensor with a volume marker in a Rnd3 knockdown cell (related to Fig. 6b (bottom)). Time-lapse TIRF video of the Rho activity sensor (delCMV-mCherry-2xRBD) and volume marker (delCMV-mCitrine). Images were collected with a frame rate of 6/min.
